# Differential microRNA expression analyses across two brain regions in Alzheimer’s disease

**DOI:** 10.1101/2021.05.31.446406

**Authors:** Valerija Dobricic, Marcel Schilling, Jessica Schulz, Ling-Shuang Zhu, Chao-Wen Zhou, Janina Fuß, Sören Franzenburg, Ling-Qiang Zhu, Laura Parkkinen, Christina M. Lill, Lars Bertram

## Abstract

**Background:** Dysregulation of microRNAs (miRNAs) is involved in the pathogenesis of neurodegenerative diseases, including Alzheimer’s disease (AD). Hitherto, sample sizes from differential miRNA expression studies in AD are exceedingly small aggravating any biological inference. To overcome this limitation, we investigated six candidate miRNAs in a large collection of brain samples.

**Methods:** Brain tissue was derived from superior temporal gyrus (STG) and entorhinal cortex (EC) from 99 AD patients and 91 controls. Expression of six miRNAs was examined by qPCR (STG) or small RNA sequencing (EC). Brain region-dependent differential miRNA expression was investigated in a transgenic AD mouse model using qPCR and FISH. Total RNA sequencing was used to assess differential expression of miRNA target genes.

**Results:** MiR-129-5p, miR-132-5p, and miR-138-5p were significantly downregulated in AD vs. controls both in STG and EC, while miR-125b-5p and miR-501-3p showed no evidence for differential expression in this dataset. In addition, miR-195-5p was significantly upregulated in EC but not STG in AD patients. The brain region-specific pattern of miR-195-5p expression was corroborated *in vivo* in transgenic AD mice. Total RNA sequencing identified several novel and functionally interesting target genes of these miRNAs involved in synaptic transmission (*GABRB1*), the immune-system response (*HCFC2*) or AD-associated differential methylation (*SLC16A3*).

**Conclusions:** Using two different methods (qPCR and small RNA-seq) in two separate brain regions in 190 individuals we more than doubled the available sample size for most miRNAs tested. Differential gene expression analyses confirm the likely involvement of miR-129-5p, miR-132-5p, miR-138-5p, and miR-195-5p in AD pathogenesis and highlight several novel potentially relevant target mRNAs.

**Funding:** This work was supported by the Deutsche Forschungsgemeinschaft (DFG) and the National Science Foundation China (NSFC) as a Joint Sino-German research project (“MiRNet-AD”, #391523883). Additional support was provided by the DFG Research Infrastructure NGS_CC (project 407495230) as part of the Next Generation Sequencing Competence Network (#423957469) and the Cure Alzheimer’ s Fund (CAF) as part of the CIRCUITS consortium project.

## Introduction

Alzheimer’s disease (AD) is the most prevalent neurodegenerative disease characterized by progressive loss of memory and cognition eventually leading to dementia. While the pathogenic mechanisms underlying AD susceptibility are not yet completely understood, it is well established that susceptibility to AD is determined by the complex interplay of genetic, environmental, and epigenetic factors. High heritability estimates both for late (LOAD) and early (EOAD) onset AD support a crucial role of genetics, but also imply the involvement of non-genetic factors. Namely, heritability for LOAD was estimated to be between 60-80% (Gatz et al., 2006), and for EOAD as >90% (Wingo et al., 2012). In addition to genetic variants, epigenetic mechanisms, e.g. mediated by DNA methylation (on the transcriptional level) and microRNAs (miRNAs; on the post-transcriptional level), are increasingly recognized to play an important role in the etiology of AD (Hébert & De Strooper, 2009; Silvestro et al., 2019; Smith et al., 2021).

MiRNAs are 18-25 nt long RNA molecules that bind to complementary sequence elements in the mRNA transcripts of protein coding genes (“target genes”) to initiate transcript degradation or translational inhibition and thus repress protein synthesis (Bartel, 2018; Guo et al., 2010). Given their important role in the regulation of gene expression, miRNAs became a topic of many studies investigating their regulatory function, role as potential biomarkers, and/or therapeutic targets for a range of human disorders, including AD (Angelucci et al., 2019; Ludwig et al., 2019). The interpretation of these studies is aggravated by various factors such as the use of heterogeneous tissues for the analysis (e.g. different brain regions, different blood cells subpopulations), application of different methods for miRNA quantification and analysis, and use of small sample sizes. Over time, this has led to a vast body of – partially contradicting – literature, which has become increasingly difficult to follow and interpret. To overcome these limitations, we recently conducted a systematic meta-analysis of differential miRNA expression studies in AD and identified 25 miRNAs showing study-wide significant differential expression in brains of AD cases vs. controls (Takousis et al., 2019).

The goal of the present study was to independently assess the top-ranking differentially expressed miRNAs in a large and carefully phenotyped collection of brain samples from AD patients and controls. Specifically, we determined the expression levels of six miRNAs in two brain regions (entorhinal cortex [EC] and superior temporal gyrus [STG]) collected from the same ∼200 individuals using either small RNA sequencing (EC) or TaqMan probe-based qPCR (STG). Moreover, brain region specific expression changes for one miRNA were assessed in different brain regions of an AD transgenic mouse model. Lastly, we probed for evidence of differential expression of mRNA targets of all analyzed miRNAs in EC using total RNA sequencing data available from the same individuals.

## Methods and Materials

### Human samples

Snap-frozen, postmortem human brain tissue from 99 AD patients and 91 elderly control individuals were obtained from the Oxford Brain Bank. These were derived from STG (Brodmann area BA21) and EC (Brodmann area BA28; for this region only n=90 AD and n=84 controls were available). The Ethics Committees of Oxford University and University of Lübeck approved the use of the human tissues for our study and all participants gave informed consent. The AD patients and healthy controls were part of the longitudinal, prospective Oxford Project to Investigate Memory and Aging (OPTIMA) using protocols which have been described in detail elsewhere (Clarke et al., 1998). All subjects underwent a detailed clinical history, physical examination, assessment of cognitive function (Cambridge Examination of Mental Disorders of the Elderly (CAMDEX) (Roth et al., 1986) with the Cambridge Cognitive Examination (CAMCOG) and Mini-Mental State Examination (MMSE) biannually. The pathological diagnosis of AD was made using the Consortium to Establish a Registry for Alzheimer’s disease (CERAD)/National Institutes of Health (NIH) criteria and Braak staging (Braak & Braak, 1991; Hyman et al., 2012; Mirra et al., 1991). All included patients were of white European descent by self report.

### Selection of miRNAs for follow-up analysis

In our recent systematic meta-analyses of differential miRNA expression studies in AD, 25 miRNAs showed study-wide (α=1.08E-04) significant differential expression in brain (Takousis et al., 2019). For the present study, we selected those showing “strong evidence” for differential expression among the top 10 miRNAs. The term “strong evidence” refers to meta-analyses with ≥80% of included studies showing the same direction of effect. In total, six miRNAs were selected: miR-125b-5p, miR-129-5p, miR-132-5p, miR-138-5p, miR-195-5p, and miR-501-3p. In addition, we ran miR-423-5p and let-7b-5p alongside to serve as endogenous controls for the qPCR assays. This selection was based on recommendations of the TaqMan Advanced miRNA assay manufacturer, previous use in the literature, and low rank or absence among AD and PD brain miRNA differential expression meta-analysis results (Schulz et al., 2019; Takousis et al., 2019).

### MiRNA isolation and quantitative PCR in superior temporal gyrus (STG) sections

For the 190 STG samples, total RNA, including miRNA, was extracted from approximately 25 mg of brain sections using the mirVana miRNA kit (Thermo Fisher Scientific, Waltham, Massachusetts) following the manufacturer’s instructions. Immediately after extraction, RNA samples were treated with DNase (TURBO DNA-free kit, Thermo Fisher Scientific). Total RNA concentration and purity were measured using a NanoDrop 2000 instrument (Thermo Fisher Scientific). Further, RNA integrity (RIN) was assessed using a Bioanalyzer 2100 instrument in conjunction with the RNA 6000 Nano LabChip kit (Agilent Technologies, Santa Clara, California).

Reverse transcription of total miRNA was carried out with the TaqMan Advanced miRNA cDNA Synthesis Kit (Thermo Fisher Scientific) using an input of 10 ng total RNA, according to manufacturer’s instructions. Quantitative assessment of the expression of miR-125b-5p (MIMAT0000423), miR-129-5p (MIMAT0000242), miR-132-5p (MIMAT0004594), miR-138-5p (MIMAT0000430), miR-195-5p (MIMAT0000461), and miR-501-3p (MIMAT0004774), along with two endogenous control miRNAs (miR-423-5p [MIMAT0004748] and let-7b-5p [MIMAT0000063]) was performed in 384-well format using TaqMan pre-spotted assays (Thermo Fisher Scientific) on a QuantStudio-12K-Flex system. Samples were assayed in triplicates. In order to minimize potential batch effects, cases and controls were randomly distributed across plates. Raw data analysis was performed using ExpressionSuite Software v1.2 (Thermo Fisher Scientific). Replicates with differences in Ct value > 0.5 to the closest replicate were excluded from subsequent analysis. All other resulting Ct values were used in the downstream differential miRNA expression analyses (see below).

### Small RNA sequencing in entorhinal cortex (EC)

EC sections were available for a subset of 174 individuals (90 cases, 84 controls) included in our current study. Quantification of the expression of the six miRNAs of interest was based on small RNA sequencing which was performed as part of another ongoing project. To this end, total RNA, including miRNA, was purified and quantified using the same methods as described above. Library preparation and subsequent sequencing were conducted at the NGS Competence Centre at IKMB institute (Kiel, Germany). Libraries were prepared using the NextFlex Small RNA-Seq kit (PerkinElmer, Waltham, Massachusetts), according to manufacturer’s instructions, and sequenced on a HiSeq 4000 instrument (Illumina, San Diego, California) with 1 × 50 bp reads. Sequencing adapters were trimmed from raw reads using Flexbar v3.4.0 (Dodt et al., 2012; Roehr et al., 2017). Reads were mapped and miRNAs quantified against miRBase v22.1 (Griffiths-Jones et al., 2006; Kozomara et al., 2019) using miRDeep2 (Friedländer et al., 2012). Data normalization was carried out using the R package DESeq2 v1.28.1 (Love et al., 2014).

### Total RNA sequencing in entorhinal cortex (EC)

Total RNA sequencing was performed at the NGS Competence Centre at IKMB (Kiel, Germany) from the same aliquots of total RNA that were also used for small RNA sequencing. Libraries were prepared using the TruSeq stranded total RNA kit (Illumina), according to manufacturer’s instructions, and sequenced on a NovaSeq 6000 instrument (Illumina) with 2 × 100 bp reads. Reads were pseudoaligned to the human transcriptome (Ensembl v. 100) (Howe et al., 2021) using kallisto v0.46.1 (Bray et al., 2016). To account for intra-sample technical variance, 100 bootstraps were performed per biological replicate. Raw reads were normalized to transcripts per million (TPM) considering protein coding isoforms of protein coding genes and lncRNA isoforms of lncRNA genes only. Isoforms with less than 6 assigned raw reads and/or less than 0.1 TPM in more than 80 % of all replicates in either condition were excluded from the analysis. Additional filtering, normalization and differential gene expression analysis was carried out using the R package sleuth (version 0.30.0) (Pimentel et al., 2017) adjusting for age at death, sex, RIN, post-mortem-interval (PMI), and the first 10 principle components of the underlying expression profiles to account for additional undetected confounding.

### Selection of putative targets of candidate miRNAs

Target predictions for miRNA families were retrieved from TargetScan v72 (McGeary et al., 2019). The predictions for the six miRNAs selected as candidates for this study were extracted by their respective seed region sequences. Predicted target mRNAs that did not qualify for differential expression analyses as per the criteria outlined above were discarded. The remaining target predictions were ranked by cumulative weighted context score (breaking potential ties by aggregate PCT and total context score). For each miRNA, the top 10 predicted target genes were selected for downstream analysis within the total RNA sequencing data. Additionally, all AD-relevant targets listed in Supplementary Table 3 of (Takousis et al., 2019) were considered, of which seven had sufficient data for differential mRNA analysis, i.e. *ADAMTS4, APP, CD2AP, CNTNAP2, FERMT2, PTK2B*, and *SORL1*.

### Statistical analysis

Log-normalized sRNA-seq counts and ΔCt values, respectively, were averaged over replicates and scaled by the corresponding mean value for let-7b-5p and miR-423-5p (endogenous controls). These values (ΔΔCt, in the case of qPCR) were transformed to relative quantity measures (2^-ΔΔCt^) and compared across conditions (AD cases vs. controls). Additionally, per miRNA and method (qPCR, sRNA-seq), a (Gaussian) generalized linear model (GLM) was fitted to predict the (scaled and centered) abundance measures from case-control status and the following (scaled and centered, if applicable) potential confounding variables: age at death, sex, RNA integrity (RIN), post-mortem interval, A260/280 absorbance. The F-statistic was used to assess the significance of the effect estimate of the AD case-control status on the expression readout. Analogously, we utilized all samples with Braak staging information to train GLMs predicting gene expression based on Braak stage. To this end, the corresponding binary (AD case vs. control) variable was replaced by a continuous (scaled and centered) Braak stage value. All other analysis steps and parameters (confounding variables) were identical to the case vs. control analyses.

For the differential miRNA expression analyses, multiple testing correction was performed using Bonferroni’s method adjusting for 6 independent miRNAs, resulting in a one-sided study-wide α of 0.0167 (=2*(0.05/6)). One-sided testing is applicable here given the specific hypotheses tested based on prior evidence from Takousis et *al*. (Takousis et al., 2019). For differential mRNA analyses, the Benjamini-Hochberg method was used to control the false discovery rate (FDR) within the 46 mRNAs assessed in this study.

### Literature search and meta-analyses

To search for novel papers on the investigated miRNAs published since the data freeze in Takousis et *al*. (Takousis et al., 2019) we performed another systematic PubMed search (www.ncbi.nlm.nih.gov/pubmed/) using the search term from the original paper “(microRNA OR miRNA OR miR OR micro-RNA) AND Alzheimer*”. We included articles published until October 1st, 2021, in peer-reviewed journals in English. Citations were assessed for eligibility using the title, abstract or the full text, as necessary. Data was extracted from studies comparing the brain expression in samples of AD patients versus controls for any of the six miRNAs analyzed here.

To arrive at summary estimates for the overall evidence of differential expression for the six tested miRNAs, we combined all data using the same meta-analysis workflow and methods as described previously (Takousis et al., 2019). Data included were 1) database assembled for Takousis et *al*., 2) novel data generated in our brain samples, and 3) novel publications on these miRNAs identified in our literature search. P-values computed for the meta-analyses represent two-sided tests, as no specific hypothesis on the effect direction was made. To correct for multiple testing, we used the same threshold as in our original publication, i.e. an α of 1.08E-04 reflecting the 461 miRNAs tested in that study (Takousis et al., 2019).

### Animal work

To confirm the region-specific differential miRNA expression of mir-195-5p observed in the human samples, we also measured brain expression patterns of miR-195-5p in EC, hippocampus and temporal cortex in two different AD mouse models (P301S and APP/PS1) through qPCR and fluorescence in situ hybridization (FISH). APP/PS1 mice (#34829) and P301S mice (#008169) were purchased from Jackson laboratory, and wild type littermates were used as control. Mice were sacrificed at 6 months of age and total RNA, including miRNA, was extracted from brain sections of control and AD mice, and reverse transcription of total miRNA was carried out as described above. After the PCR reaction, amplified DNA fragments were verified by gel electrophoresis on a 3% agarose gel. Amplification and analysis were performed in the iCycler iQ Multicolor Real-Time PCR Detection System (BioRad, Hercules, California).

After loading on HistoBond Slides (VWR, Radnor, Pennsylvania) and drying overnight at 42 °C, 30 μm brain slices were incubated in protease K detergent at 37 °C for 30 min, and then transferred to 0.25% acetic anhydride and 0.1M triethanolamine for 10 min. Hybridization buffers containing 5’ HEX-labeled probes were used to incubate slices in an in-situ hybridization apparatus (Boekel Slide Moat, Feasterville, Pennsylvania) at 52 °C under the condition of heat preservation and humidification. Subsequently, brain slices were incubated with anti-Calbindin D-28k (Swant, #300, diluted in 1:500) and Alexa Flour 546 (ThermoFisher, diluted in 1:200). The images were collected using a confocal laser scanning microscope (Axio Imager Z2,[Zeiss, Oberkochen, Germany], Motorized Scanning Stage [Maerzhaeuser, Wetzlar, Germany]), and analyzed using Zen Pro. Probes were designed and purchased from Tsingke, China.

All animal experiments were performed according to the “Policies on the Use of Animals and Humans in Neuroscience Research” revised and approved by the Society for Neuroscience in 1995. The conduct of all animal experiments was approved by the Animal Ethics Committee of Huazhong University of Science and Technology.

## Results

### Demographics and RNA quality assessments

The average age of death was 81.59 years (median = 83 years, interquartile range [IQR] = 9.5 years) for AD patients and 77.48 years (median = 81 years, IQR = 20 years) for controls (Welch t-test, P = 0.0142). The average postmortem interval was 57.07h (median = 48 h, IQR = 43.75 h) for AD cases, and 48.41 h (median = 48 h, IQR = 24 h) for controls (Welch t-test, P = 0.054) (Table 1). There was no significant difference in the sex distribution between the AD and the control group (chi-square test, P = 0.536). Detailed distributions of Braak stages are given in Table 1. Numbers above are for the larger STG dataset (n=190) but were equivalent for the EC dataset (n=174).

**Table 1.**
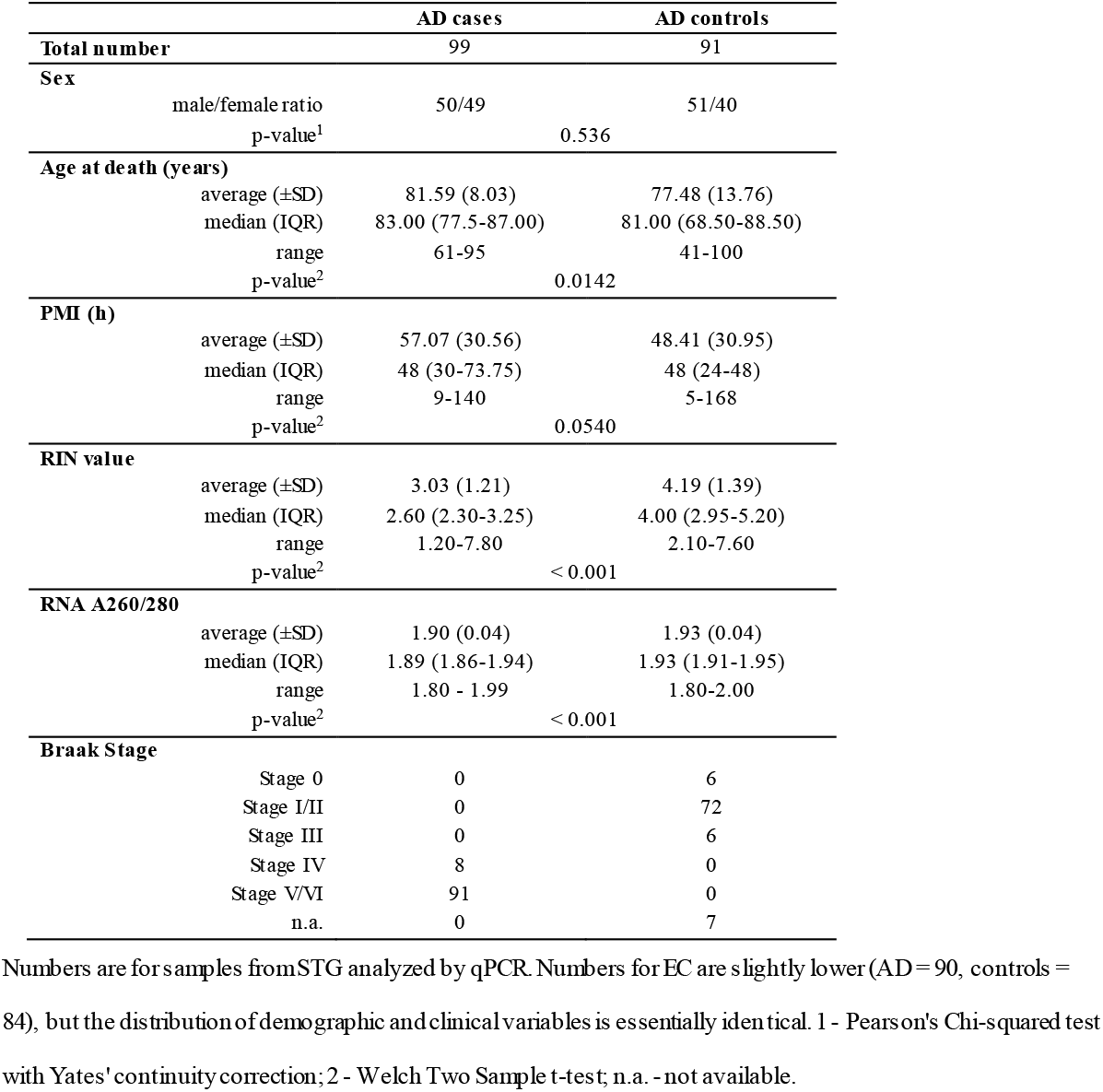
Overview of the brain samples analyzed in this study.

|  | AD cases | AD controls |
| --- | --- | --- |
| **Total number** | 99 | 91 |
| **Sex** |  |  |
| male/female ratio | 50/49 | 51/40 |
| p-value<sup>1</sup> | 0.536 |  |
| **Age at death (years)** |  |  |
| average (±SD) | 81.59 (8.03) | 77.48 (13.76) |
| median (IQR) | 83.00 (77.5-87.00) | 81.00 (68.50-88.50) |
| range | 61-95 | 41-100 |
| p-value<sup>2</sup> | 0.0142 |  |
| **PMI (h)** |  |  |
| average (±SD) | 57.07 (30.56) | 48.41 (30.95) |
| median (IQR) | 48 (30-73.75) | 48 (24-48) |
| range | 9-140 | 5-168 |
| p-value<sup>2</sup> | 0.0540 |  |
| **RIN value** |  |  |
| average (±SD) | 3.03 (1.21) | 4.19 (1.39) |
| median (IQR) | 2.60 (2.30-3.25) | 4.00 (2.95-5.20) |
| range | 1.20-7.80 | 2.10-7.60 |
| p-value<sup>2</sup> | < 0.001 |  |
| **RNA A260/280** |  |  |
| average (±SD) | 1.90 (0.04) | 1.93 (0.04) |
| median (IQR) | 1.89 (1.86-1.94) | 1.93 (1.91-1.95) |
| range | 1.80 - 1.99 | 1.80-2.00 |
| p-value<sup>2</sup> | < 0.001 |  |
| **Braak Stage** |  |  |
| Stage 0 | 0 | 6 |
| Stage I/II | 0 | 72 |
| Stage III | 0 | 6 |
| Stage IV | 8 | 0 |
| Stage V/VI | 91 | 0 |
| n.a. | 0 | 7 |

Numbers are for samples from STG analyzed by qPCR. Numbers for EC are slightly lower (AD = 90, controls = 84), but the distribution of demographic and clinical variables is essentially identical. 1 - Pearson's Chi-squared test with Yates' continuity correction; 2 - Welch Two Sample t-test; n.a. - not available.

RNA yield was good across all STG (EC) samples with 490 ng (650 ng) total RNA per mg of tissue weight. RIN values ranged between 1.2 (1.2) and 7.8 (6.3), with an average of 3.6 (3.0). Comparison of raw expression data showed that the distribution of Ct values (across all miRNA assays per sample) was similar for samples with lower (RIN < 5) vs. higher RIN values (RIN ≥ 5) (Supplementary Figure 1). The same was observed for the distribution of average Ct values for endogenous control assays in the samples with lower (RIN < 5) and higher RIN values (RIN > 5) (mean Ct_RIN<5_ = 23.92 vs. mean Ct_RIN ≥5_= 23.99), indicating that miRNAs were not majorly affected by RNA degradation (data not shown), in line with Lau et *al*. (Lau et al., 2013). Notwithstanding, we adjusted for differences in average RIN and absorbance values in AD samples vs. controls (RIN_AD_ =3.03 vs. RIN_ctrl_ = 4.19; Welch t-test, P<0.001; A260/280_AD_ = 1.90 vs. A260/280_ctrl_ = 1.93, Welch t-test, P<0.001) in the regression models to account for residual confounding.

### Differential miRNA expression analysis in brains of AD cases and controls

Our qPCR-based expression analyses showed that three (i.e. miR-132-5p [P = 1.80E-21], miR-138-5p [P = 2.80E-04], miR-129-5p [P = 1.50E-08]) of the six tested miRNAs, showed evidence for significant differential expression in STG sections in AD cases vs. controls (Table 2). All three miRNAs showed *decreased* expression levels in AD cases. Likewise, in EC, a significant downregulation of the same three miRNAs in AD patients was observed using a different experimental method (i.e. small RNA sequencing). In addition, we observed miR-195-5p to be significantly upregulated in EC of AD vs. controls (P = 8.40E-05), but not in STG (P = 0.088). We observed no evidence for differential expression for neither of the other two tested miRNAs in either STG or in EC (Figure 1, Table 2). With the exception of miR-125b-5p and miR-501-3p, the direction of expression change was concordant in STG vs. EC for the remaining four miRNAs (Table 2).

**Table 2.** Results of targeted miRNA differential expression analysis in two different brain regions.

| miR name<br>(miRBASE) | Current study, STG |  |  |  | Current study, EC |  |  |  | Meta-analysis (Takousis *et al.*) |  |  |  | Current study, Meta-analysis |  |  |  |
| --- | --- | --- | --- | --- | --- | --- | --- | --- | --- | --- | --- | --- | --- | --- | --- | --- |
|  | Direction | P-value<sup>§</sup> | Effect size<br>(±SE) | N (cases,<br>ctrls) | Direction | P-value<sup>§</sup> | Log2 Fold<br>Change (SE) | N (cases,<br>ctrls) | Direction | P-value<sup>#</sup> | N (cases,<br>ctrls) | N<br>studies | Direction | P-value<sup>#</sup> | N (cases,<br>ctrls) | N<br>studies |
| hsa-miR-125b-5p | down | 0.605 | -0.0742<br>(±0.143) | 190 (99, 91) | up | 0.233 | 0.059<br>(±0.0547) | 174 (90, 84) | up | **4.13E-13** | 122 (64, 58) | 11 | up | **4.15E-05** | 312 (163,<br>149) | 12 |
| hsa-miR-501-3p | down | 0.902 | -0.0211<br>(±0.171) | 174 (89, 85) | up | 0.0465 | 0.139<br>(±0.0891) | 174 (90, 84) | up | **2.03E-11** | 68 (38, 30) | 4 | up | **1.56E-06** | 308 (154,<br>154) | 6 |
| hsa-miR-132-5p | down | **1.75E-21** | -1.18<br>(±0.109) | 190 (99, 91) | down | **1.15E-04** | -0.395<br>(±0.131) | 174 (90, 84) | down | **3.01E-10** | 57 (27, 30) | 5 | down | **3.67E-35** | 313 (153,<br>160) | 7 |
| hsa-miR-138-5p | down | **2.82E-04** | -0.525<br>(±0.142) | 190 (99, 91) | down | **2.46E-04** | -0.362<br>(±0.118) | 174 (90, 84) | down | **5.50E-08** | 161 (96, 65) | 6 | down | **2.40E-10** | 351 (195,<br>156) | 7 |
| hsa-miR-195-5p | up | 0.088 | 0.260<br>(±0.152) | 190 (99, 91) | up | **8.41E-05** | 0.295<br>(±0.0856) | 174 (90, 84) | up | **3.74E-07** | 177 (104, 73) | 7 | up | **2.29E-07\*** | 433 (230,<br>203) | 9 |
| hsa-miR-129-5p | down | **1.54E-08** | -0.705<br>(±0.119) | 190 (99, 91) | down | **7.00E-09** | -0.705<br>(±0.135) | 174 (90, 84) | down | **3.84E-07** | 166 (100, 66) | 6 | down | **6.25E-16** | 1122 (653,<br>469) | 9 |

§ - one-sided p-values; # - two-sided p-values; cases – Alzheimer's disease patients; Ctrl's - controls; bold front - differential expression reaching significance
threshold for the corresponding analysis ( $\alpha$ (one-sided)=0.0167 for differential expression analyses, $\alpha$ (two-sided)=1.08E-04 for meta-analyses; see Methods);
\*The equivalent P-value after meta-analysis using EC instead of STG data is 2.39E-11; SE - standard error.

**Figure 1.**
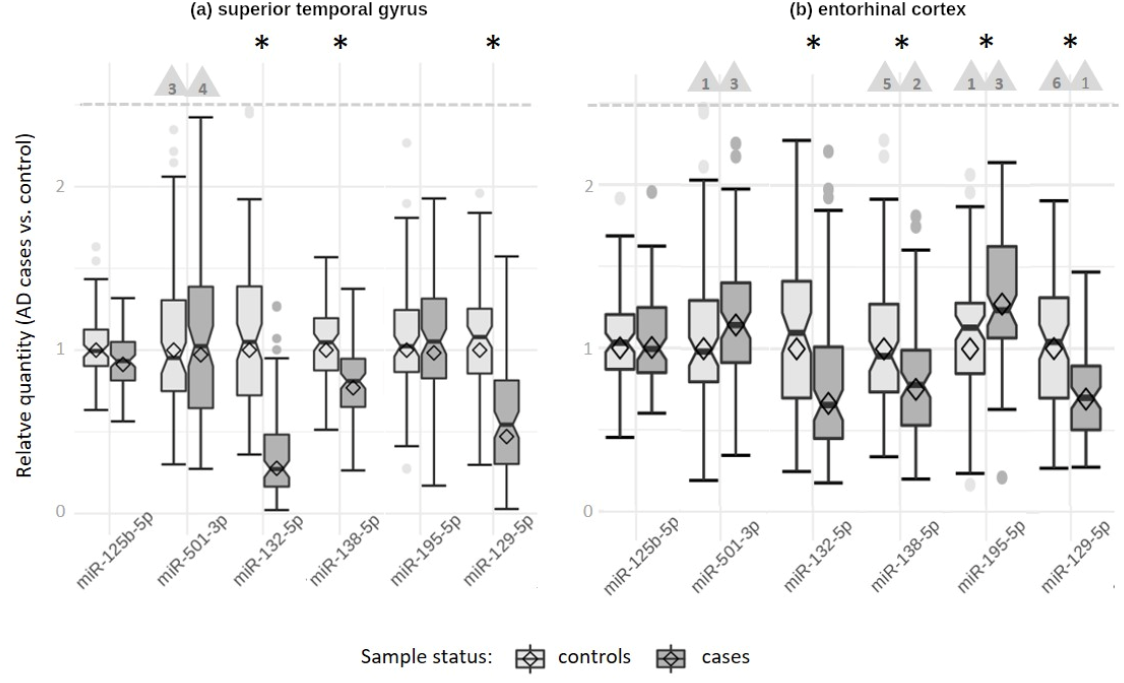
Expression levels of analyzed miRNAs in Alzheimer’s disease patients relative to controls. <u>Panel A</u>: Superior temporal gyrus (STR) samples analyzed by qPCR. <u>Panel B</u>: Entorhinal cortex (EC) samples analyzed by small RNA sequencing. Bars filled in light grey: controls; bars filled in dark grey: AD cases; *: statistically significant difference at a = 0.0167 (see Methods). The relative quantity of miRNA expression was calculated using the ΔΔCt method; diamonds represent the mean expression (cases relative to controls) based on the ΔΔCt method (relative quantity = 2(-(dCt cases - dCt controls))). Horizontal lines represent median values of the corresponding sample-specific values (individual dCt values normalized to the mean of the control samples), boxes represent interquartile ranges, and whiskers extend to the minimum and maximum observed value within 1.5x the interquartile range; values outside this range but below the dashed line are depicted as dots. The box notches indicate the 95 % confidence intervals. Outliers exceeding the dashed line are not shown (for scaling purposes) but counted and indicated by the numbers in the triangles.

Using Braak-staging as diagnostic variable yielded similar results to those obtained in the case vs. control analyses (Supplementary Table 1). In STG, these highlighted miR-129-5p, miR-138-5p, and miR-132-5p as differentially expressed. In EC, these analyses revealed the same three miRNAs and miR-195-5p.

### Assessment of brain region-specific expression changes for miR-195-5p in two different AD mouse models

We further examined the expression of miR-195-5p in different brain regions of 6-month-old APP/PS1 and P301S mice to assess whether the region-specific expression difference for this miRNA observed in the human samples can be recapitulated in these models. Consistent with the results observed in the human samples, we observed an upregulation of miR-195-5p in EC of P301S mice vs controls (p=0.0226), but not in hippocampus and temporal cortex (Figure 2). No differences were found in any of the analyzed brain regions of APP/PS1 mice vs. controls, indicating that miR-195-5p upregulation might be related to tau pathology, for which P301S is a model. Furthermore, FISH analysis showed that miR-195-5p is mainly increased in EC layer II Calb+ neurons, the most vulnerable neurons in the early stage of AD (Yassa, 2014), in P301S but not APP/PS1 mice (Figure 2).

**Figure 2.**
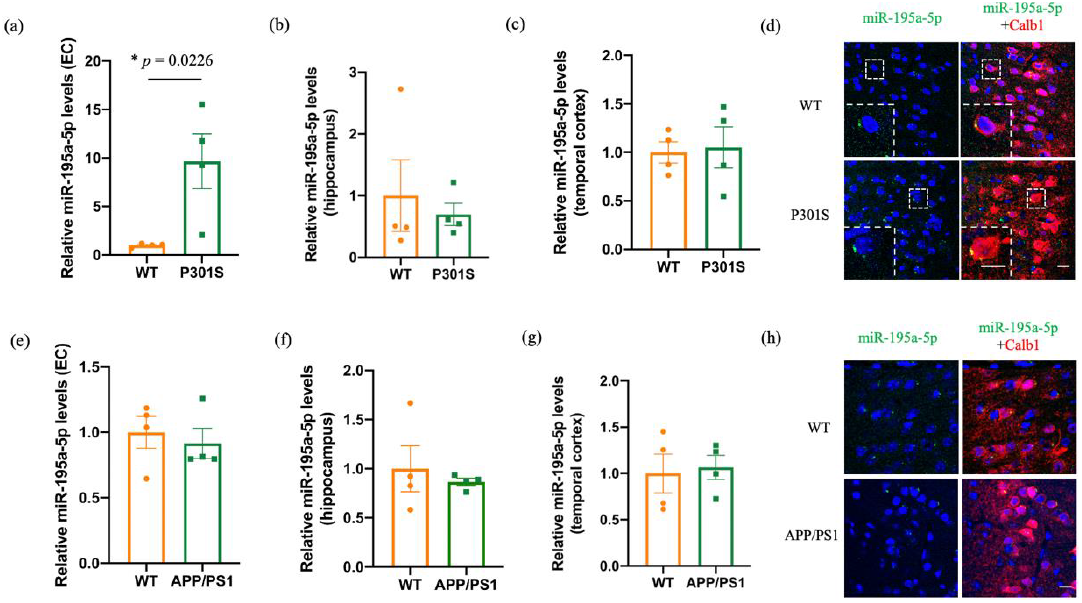
Results of qPCR and fluorescence in situ hybridization (FISH) experiments on miR-195a-5p expression in two different AD mouse models compared to controls. <u>Panel A</u>: Expression of miR-195a-5p in entorhinal cortex of 6-month-old P301S mice compared to wild-type (WT) mice was assessed by qPCR (n = 4 for each group). * p-value from unpaired Student’s t test was used comparing transgenic vs. WT mice. <u>Panels B-C</u>: Expression of miR-195a-5p in hippocampus and temporal cortex of 6-month-old P301S mice compared to WT mice; no significant differences were identified. <u>Panel D</u>: Fluorescence in-situ hybridization (FISH) images of miR-195a-5p which is stained green in EC of 6-month-old P301S and WT mice. The nuclei were stained blue with DAPI. Cabl1 (stained in red) was used as a marker of EC layer II. Scale bar indicates 20 μm. n = 3 mice in each group. <u>Panels E-H</u>: Analogous experiments comparing EC (e), hippocampus (f) and temporal (g) cortex and performing FISH staining (h) in 6-month-old APP/PS1 mice compared to WT mice.

### Meta-analysis of novel differential miRNA expression results with published evidence

To assess the overall evidence of differential expression of the six miRNAs tested here we updated our earlier meta-analyses by combining 1) results from the database of Takousis et *al*., 2) STG-based results from the current study, and 3) data from additional studies testing any of the six miRNAs for differential miRNA expression in human brain samples not included in our previous report (Li & Cai, 2021; Patrick et al., 2017).

All updated meta-analyses are shown in Table 2. There are at least three noteworthy observations to be made from these results: First, for all three miRNAs significant in our own brain dataset (i.e. miR-132-5p, miR-138-5p, and miR-129-5p; see above), the statistical evidence increased by several orders of magnitude (2-25x, as judged by P-value) upon meta-analysis. Second, in contrast to these strengthened results, two miRNAs that ranked very high in our previous assessment of the literature (Takousis et al., 2019), i.e. miR-125b-5p and miR-501-3p, were not replicated in our independent brain dataset and, as a result, now only show reduced overall evidence of differential expression upon meta-analysis. We note that of these two miRNAs only miR-501-3p was studied in the newly included paper by Li & Cai, 2021, who found a significant up-regulation in AD vs. controls (and in line with the previous meta-analysis). Notwithstanding, the overall meta-analytic evidence combining all newly available data for this miRNA showed reduced statistical support (P-value 1.56E-06) when compared to our previous study (P-value 2.03E-11). Third, for one miRNA (i.e. miR-195-5p) we observed region-specific differences in differential expression (significant only in EC; see above). As a result, only the meta-analyses using data from EC (p=2.39E-11; Supplementary Table 1) improved with respect to those from our previous study (p=3.74E-07; (Takousis et al., 2019)). Taken together, our novel differential miRNA expression results derived from a large and independent dataset analyzed in combination with previously published data now nominate a revised and partially different set of miRNAs to be most strongly linked to AD in brain.

### Differential miRNA target expression assessment in matching totalRNA-seq data

Lastly, we assessed whether and which target genes of the miRNAs tested in our primary smallRNA-seq analysis arm also show evidence for differential expression in the same individuals. MiRNA target predictions were taken from the TargetScan database, which returned informative mRNA predictions except for hsa-miR-132-5p (family/seed: CCGUGGC). Considering only the top 10 targets (see Methods) for the remaining five candidate miRNAs as well as additional AD-related targets reported in (Takousis et al., 2019) resulted in 53 unique mRNAs (five genes were among the selected targets of two different candidate miRNAs). Five out of these showed evidence for nominally significant differential expression (p<0.05) according to our total RNA-seq data. We note, however, that none of these nominally significant results attained study-wide significance after adjusting for multiple testing across all 53 tests (pFDR < 5%). Differential gene expression results are listed in Supplementary Table 3.

Interestingly, for several of the top (ranked by p-value) differentially expressed mRNAs previous studies had already implicated a potential functional link to AD. Among these are *SLC16A3, HCFC2*, and *GABRB1*. Gene *SLC16A3* (p-value 0.00347; protein: solute carrier family 16 member 3) is a member of the proton-linked monocarboxylate transporter (MCT) family and is involved in the transport of lactic acid and pyruvate across plasma membranes (https://www.ncbi.nlm.nih.gov/nuccore/1890259789). This gene has been linked to AD by several independent epigenome-wide association studies (EWAS) where it showed significant association with either Braak-stage (Smith et al., 2021) or AD diagnostic status (Li et al., 2020) in human brain samples. Other work recently implied *SLC16A3* and other members of the MCT family to show differential expression in AD oligodendrocytes in human brain (Saito et al., 2021). *HCFC2* (p-value 0.0395; protein: host cell factor C2) encodes one of two proteins which interact with VP16, a herpes simplex virus protein that initiates virus infection (https://www.ncbi.nlm.nih.gov/nuccore/NM_013320). A potential link to AD pathogenesis was recently implied by weighted gene co-expression network analysis suggesting that *HCFC2* is one of several factors involved in differential immune cell infiltration in AD prefrontal cortex (Liu et al., 2022). Lastly, *GABRB1* (p-value 0.0534; protein: gamma-aminobutyric acid type A receptor subunit beta1) encodes a subunit of the GABA-A neurotransmitter receptor that mediates inhibitory synaptic transmission in the central nervous system (https://www.ncbi.nlm.nih.gov/nuccore/NM_000812). In addition, recent *in vivo* work using transgenic mouse models suggests that GABRB1 may serve as a synaptic receptor for secreted APP (sAPP) providing functional support of the long-sought link between sAPP and synaptic transmission (Rice et al., 2019). We note that we restricted our miRNA target gene analyses to only the top10 targets provided on TargetScan. As a result, this arm of our study is – by design – incomplete and should only be understood as a first exemplary discussion of the potential functional implications of our primary miRNA differential expression experiments.

In addition to the target genes predicted by the TargetScan database, we also investigated whether any of the AD-relevant (i.e. implicated by GWAS) target genes highlighted in our previous work (Takousis et al., 2019) also showed evidence for differential expression in the EC brain sections analyzed here (genes marked by *; Supplementary Table 3). While none of the seven tested candidate targets passed the threshold of nominal significance, we note that two (i.e. *ADAMTS4* and *CNTNAP2*) approached significance with p-values of 0.0502 and 0.0687, respectively. All of the remaining five tested AD candidates (i.e. *APP, CD2AP, FERMT2, PTK2B*, and *SORL1*) showed differential gene expression p-values >0.1.

## Discussion

In this study, we performed well-powered and independent assessments of the most compelling miRNAs previously reported to show differential expression in brains of AD patients. Specifically, we analyzed the expression of six “top” miRNAs from these meta-analyses in two separate cortical regions in a comparatively large (n∼200) dataset using two experimental methods of miRNA quantification. The results showed evidence for significant differential expression for three out of six miRNAs tested in STG and four out of six in EC. One AD miRNA (miR-195-5p) showed brain region-specific differential miRNA expression, i.e. a significant upregulation in AD in EC but not STG slices from the same individuals, a finding that was corroborated in an AD transgenic mouse model. We updated our previous meta-analyses on these miRNAs with the novel data collected here. The results now nominate a partially different set of top miRNAs to be linked to AD in brain, with miR-132-5p, miR-129-5p, miR-138-5p, and miR-195-5p now representing the most strongly implicated miRNA candidates. Lastly, we performed differential mRNA expression analyses on target genes of dysregulated miRNAs using total RNA-seq data generated in the same individuals. These analyses revealed a number of differentially expressed genes, some of which have previously reported links to AD or AD-relevant traits emphasizing the likely functional relevance of our findings.

Our novel results are noteworthy for several reasons. First, by analyzing brain sections from nearly 200 individuals, our study vastly increases the total available sample size for all six miRNAs tested. This is important given the observation that the median sample size of miRNA differential expression studies in brain in AD was only 42.5 (inter quartile range [IQR] 23-85) (Takousis et al., 2019) prior to this study. Second, by analyzing two different brain regions (EC and STG) using two different experimental methods (qPCR and small-RNA sequencing) our results are relatively well protected against tissue- or methods-related artifacts. This is further evidenced by the fact that most (but not all, see below) differential miRNA expression results correspond well to the prior evidence. Third, by updating the previous meta-analyses from Takousis et al. with both our novel data as well as other data from study published since our original assessment, the results provided herein represent the most current snapshot of miRNA expression data in the field. Specifically, the new meta-analyses considerably strengthened the evidence for four of the six tested miRNAs, i.e. miR-129-5p, miR-132-5p, miR-138-5p, and miR-195-5p. Interestingly, for the latter, we only observed significant differential expression in EC but not in STG in both human and mouse data, arguing for the need to analyze multiple brain regions in future studies. In contrast, two miRNAs previously showing very strong evidence for differential expression in AD, i.e., miR-125b-5p and miR-501-3p, are no longer on the top of the list. Their drastic drop in significance (and perhaps functional importance) is the result of the size of our dataset, which exceeds the previously published sample sizes for these miRNAs by several-fold, respectively, in nearly all cases. While it remains possible that the non-validation in our dataset reflects a false-negative finding, this appears unlikely given the consistency of our null findings across both brain regions and both molecular methods used. Hence, our data suggest that miR-125b-5p and miR-501-3p may be less relevant in AD pathogenesis than previously thought. Finally, using differential mRNA expression data derived from total RNA sequencing experiments performed in the same individuals, we identified several functionally interesting target genes of the top differentially expressed miRNAs. AD-relevant functional domains affected include synaptic transmission (*GABRB1*) and potentially the immune-system response (*HCFC2*). While for *SLC16A3*, which showed the strongest evidence of differential target gene expression in our study, no direct functional connection to AD has been made to date to our knowledge, this gene was recently associated with AD by several brain-based EWAS. Despite the interesting functional candidacy of these and other loci in our list of miRNA target genes (Supplementary Table 3), we note that the differential expression evidence for any of these genes was only significant at a nominal level. Future work in larger data sets needs to assess the role of these and other target mRNAs in AD pathophysiology.

Despite its strengths, our study may also be subject to a number of limitations. First, while our sample size (n∼200) was large compared to most previous studies on the topic (medium sample size = 42.5; largest previous meta-analysis sample size for the miRNAs studied = 177; (Takousis et al., 2019)), it may have still been too small to detect minor differences in miRNA expression, so that all or some of our null findings may reflect false negatives. Second, with an average of 3.6 the RIN values of our samples was comparatively low, which may have led to both false positive as well as false-negative results. However, we went to great lengths at accounting for this limitation in our analyses (see methods and results) and found no evidence that low RIN values actually skewed our differential miRNA expression results. Moreover, there are multiple studies reporting that RIN values only had a negligible or no effect on the detection of miRNAs, unlike mRNAs which tend to gradually degrade with decreasing RIN values (Jung et al., 2010). In addition, the fact that most of the previous “top” miRNAs actually do show independent replication here also argues against a major impact of low RIN on our study. Notwithstanding, we cannot exclude the possibility that the comparatively small RIN values may have affected all or some of the mRNA differential expression analyses. Third, despite being comparatively comprehensive in both size and scope, our study used RNA extracts from “bulk” brain sections. These comprise a mixture of different cell-types (e.g. neurons, immune cells) which may have confounded some of our results. The only *bona fide* remedy against this potential confounding would be to perform single-cell/single-nucleus RNA sequencing. However, this methodology is currently still comparatively expensive precluding analyses in sample sizes such as achieved here in the foreseeable future.

Since our study followed-up on previous work, the potential functional implications of the miRNAs highlighted to show consistent and highly significant differential expression here have not changed much and we refer to the discussion of Takousis et *al*. for more details. The most interesting aspect in this context is probably the assessment of whether or not the four validated miRNAs of this study target any of the known AD genes as judged by the 2013 GWAS from the IGAP (Lambert et al., 2013). In the Takousis *et al*. report this had revealed a total of seven AD genes for the four miRNAs validated in our study, i.e. *ADAMTS4* (miR-129-5p), *APP* (miR-138-5p and miR-195-5p), *CD2AP* (miR-195-5p), *CNTNAP2* (miR-195-5p), and *FERMT2* (miR-138-5p). Comparing the same target predictions to an updated list of AD genes identified by from two more recent GWAS (Jansen et al., 2019; Kunkle et al., 2019), as summarized in Bertram & Tanzi (Bertram & Tanzi, 2020) did not change these predictions. However, using an extended and even more recent list of GWAS results from the European Alzheimer’s disease DNA biobank (EADB) project published as preprint (Bellenguez et al., 2020) reveals several new connections, i.e. for *ADAM17 & USP6NL* (both miR-129-5p), *CTSB & EED* (miR-138-5p), and *ANK3 & PLEKHA1* (miR-195-5p). Collectively, these results – and those from the mRNA differential expression analyses newly performed in our EC brain slices – offer a direct link between two different molecular layers both showing an involvement in AD pathogenesis using entirely different methodologies. As such, they provide some first functional leads on the potential mechanisms by which the miRNAs found to be differentially expressed in our and previous work may unfold their effects. We note, however, that none of the AD-relevant target genes highlighted above showed strong evidence for differential mRNA expression in our dataset, so that future work is needed to further assess these potential functional implications.

In conclusion, by studying the expression patterns of six previously top-ranked miRNAs across two human brain regions in a sample of ∼200 AD patients and control individuals, we confirm the likely involvement of miR-129-5p, miR-132-5p, miR-138-5p, and miR-195-5p in AD pathogenesis.

## Acknowledgements

This work was supported by the Deutsche Forschungsgemeinschaft (DFG) and the National Science Foundation China (NSFC) as a Joint Sino-German research project (“MiRNet-AD”, #391523883) to L.B. and L.-Q.Z. Additional support was provided by the DFG Research Infrastructure NGS_CC (project 407495230) as part of the Next Generation Sequencing Competence Network (#423957469) and the Cure Alzheimer’s Fund (as part of the CIRCUITS consortium). NGS analyses were carried out at the Competence Centre for Genomic Analysis (Kiel). We acknowledge the Oxford Brain Bank, supported by the Medical Research Council (MRC), Brains for Dementia Research (BDR) (Alzheimer Society and Alzheimer Research UK), Autistica UK and the NIHR Oxford Biomedical Research Centre.

