## Supplementary for "Differential microRNA expression analyses across two brain regions in Alzheimer’s disease"

**Supplementary Figure 1**. Box plot displaying the distribution of qPCR-based Ct values for each sample analyzed in this study.

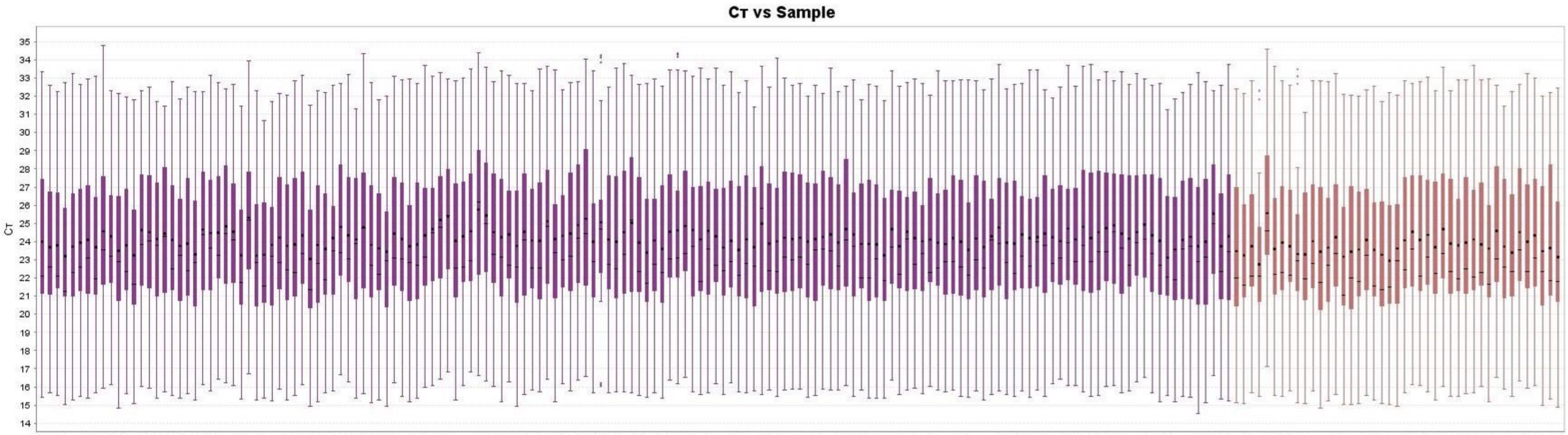


Samples (on x-axis) with RIN<5 are represented in purple and samples with RIN ≥5 in brown color. The solid box shows the range of the middle 50% of the Ct values for each sample. The horizontal black bar shows the median Ct value. The black circle shows the mean Ct value.

**Supplementary Table 1**. Results of targeted miRNA differential expression analysis in two different brain regions using Braak staging as diagnostic variable.

| miR name (miRBASE) | Current study, STG | | | | | Current study, EC | | | | |
|---|---|---|---|---|---|---|---|---|---|---|
| | Direction | P-value | Effect size (±SE) | CI | N (cases, ctrls) | Direction | P-value | Effect size (±SE) | CI | N (cases, ctrls) |
| hsa-miR-125b-5p | down | 0.301 | 0.0756 (±0.0729) | -0.218, 0.0672 | 190 (99, 91) | up | 0.496 | 0.0465 (±0.0682) | -0.087, 0.180 | 174 (90, 84) |
| hsa-miR-501-3p | down | 0.950 | -0.00547 (±0.0874) | -0.177, 0.166 | 174 (89, 85) | up | 0.0321 | 0.0804 (±0.0372) | 0.00751, 0.153 | 174 (90, 84) |
| hsa-miR-132-5p | down | **2.76E-23** | -0.620 (±0.0538) | -0.726, -0.515 | 190 (99, 91) | down | **1.12E-05** | -0.354 (±0.078) | -0.507, -0.201 | 174 (90, 84) |
| hsa-miR-138-5p | down | **3.30E-05** | -0.305 (±0.0716) | -0.446, -0.165 | 190 (99, 91) | down | **1.10E-04** | -0.0965 (±0.0243) | -0.144, -0.0488 | 174 (90, 84) |
| hsa-miR-195-5p | up | 0.169 | 0.105 (±0.0757) | -0.0438, 0.253 | 190 (99, 91) | up | **1.30E-04** | 0.191 (±0.0488) | 0.0958, 0.287 | 174 (90, 84) |
| hsa-miR-129-5p | down | **9.89E-12** | -0.422 (±0.0579) | -0.536, -0.309 | 190 (99, 91) | down | **1.41E-04** | -0.248 (±0.0636) | -0.373, -0.123 | 174 (90, 84) |

Cases – Alzheimer's disease patients; ctrls - controls; bold font - differential expression reaching significance threshold after multiple testing correction (one-sided test, $\alpha = 0.0167$; see Methods); SE - standard error; CI - confidence interval.

**Supplementary Table 2**. Results of meta-analyses combining data from Takousis et *al*., new publications identified in our literature search, and the novel data generated in EC brain samples of our project (for meta-analysis results using STG data see Table 2).

| miR name (miRBASE) | Direction | P-value | Cases | Ctrls | N total | N studies |
|---|---|---|---|---|---|---|
| hsa-miR-125b-5p | up | **2.80E-08** | 154 | 142 | 296 | 12 |
| hsa-miR-501-3p | up | **1.46E-10** | 155 | 153 | 308 | 6 |
| hsa-miR-132-5p | down | **9.38E-16** | 144 | 153 | 297 | 7 |
| hsa-miR-138-5p | down | **1.68E-10** | 186 | 149 | 335 | 7 |
| hsa-miR-195-5p | up | **2.39E-11** | 221 | 196 | 417 | 9 |
| hsa-miR-129-5p | down | **5.74E-16** | 644 | 462 | 1106 | 9 |

Cases – Alzheimer's disease patients; ctrls - controls; bold font - differential expression reaching significance threshold (two-sided test, $\alpha = 1.08E-04$; see Methods);

**Supplementary Table 3**. Results of targeted mRNA differential expression analysis on top predicted target genes of miRNAs reported in Table 2.

| Ensembl gene ID | Target gene name | miRNA(s) predicted to bind target gene‡ | FC | P-value | Q-value |
|---|---|---|---|---|---|
| **ENSG00000141526** | ***SLC16A3*** | **hsa-miR-129-5p** | **0.954** | **0.00347** | **0.184** |
| **ENSG00000132002** | ***DNAJB1*** | **hsa-miR-125b-5p** | **0.884** | **0.0323** | **0.404** |
| **ENSG00000111727** | ***HCFC2*** | **hsa-miR-195-5p** | **1.11** | **0.0395** | **0.404** |
| **ENSG00000127083** | ***OMD*** | **hsa-miR-129-5p** | **1.77** | **0.043** | **0.404** |
| **ENSG00000168938** | ***PPIC*** | **hsa-miR-129-5p** | **1.50** | **0.05** | **0.404** |
| ENSG00000158859 | *ADAMTS4** | hsa-miR-129-5p | 0.941 | 0.0502 | 0.404 |
| ENSG00000163288 | *GABRB1* | hsa-miR-138-5p | 0.711 | 0.0534 | 0.404 |
| ENSG00000174469 | *CNTNAP2** | hsa-miR-195-5p | 0.554 | 0.0687 | 0.416 |
| ENSG00000163935 | *SFMBT1* | hsa-miR-138-5p | 0.755 | 0.0706 | 0.416 |
| ENSG00000166006 | *KCNC2* | hsa-miR-125b-5p | 0.541 | 0.0821 | 0.435 |
| ENSG00000181722 | *ZBTB20* | hsa-miR-138-5p, hsa-miR-501-3p | 1.47 | 0.0908 | 0.437 |
| ENSG00000070193 | *FGF10* | hsa-miR-138-5p | 0.97 | 0.131 | 0.478 |
| ENSG00000196862 | *RGPD4* | hsa-miR-195-5p | 0.734 | 0.136 | 0.478 |
| ENSG00000105135 | *ILVBL* | hsa-miR-501-3p | 0.965 | 0.141 | 0.478 |
| ENSG00000073712 | *FERMT2** | hsa-miR-138-5p | 0.985 | 0.143 | 0.478 |
| ENSG00000125520 | *SLC2A4RG* | hsa-miR-129-5p | 1.03 | 0.144 | 0.478 |
| ENSG00000235568 | *NFAM1* | hsa-miR-195-5p | 1.13 | 0.175 | 0.528 |
| ENSG00000177181 | *RIMKLA* | hsa-miR-138-5p | 0.615 | 0.179 | 0.528 |

| | | | | | |
|---|---|---|---|---|---|
| ENSG00000137642 | *SORL1** | hsa-miR-125b-5p | 0.899 | 0.206 | 0.558 |
| ENSG00000176407 | *KCMF1* | hsa-miR-501-3p | 0.832 | 0.221 | 0.558 |
| ENSG00000168291 | *PDHB* | hsa-miR-129-5p, hsa-miR-195-5p | 0.757 | 0.221 | 0.558 |
| ENSG00000138443 | *ABI2* | hsa-miR-501-3p | 0.809 | 0.261 | 0.601 |
| ENSG00000143473 | *KCNH1* | hsa-miR-125b-5p, hsa-miR-138-5p | 0.622 | 0.271 | 0.601 |
| ENSG00000169306 | *IL1RAPL1* | hsa-miR-125b-5p | 1.09 | 0.272 | 0.601 |
| ENSG00000124249 | *KCNK15* | hsa-miR-138-5p, hsa-miR-195-5p | 0.798 | 0.286 | 0.605 |
| ENSG00000198087 | *CD2AP** | hsa-miR-195-5p | 1.24 | 0.305 | 0.607 |
| ENSG00000118473 | *SGIP1* | hsa-miR-195-5p | 0.679 | 0.314 | 0.607 |
| ENSG00000068489 | *PRR11* | hsa-miR-195-5p | 0.927 | 0.321 | 0.607 |
| ENSG00000144406 | *UNC80* | hsa-miR-501-3p | 0.663 | 0.349 | 0.637 |
| ENSG00000197548 | *ATG7* | hsa-miR-138-5p | 1.05 | 0.394 | 0.671 |
| ENSG00000182985 | *CADM1* | hsa-miR-501-3p | 0.946 | 0.405 | 0.671 |
| ENSG00000169629 | *RGPD8* | hsa-miR-195-5p | 0.923 | 0.418 | 0.671 |
| ENSG00000155052 | *CNTNAP5* | hsa-miR-125b-5p | 0.551 | 0.424 | 0.671 |
| ENSG00000120675 | *DNAJC15* | hsa-miR-138-5p | 1.14 | 0.456 | 0.671 |
| ENSG00000183454 | *GRIN2A* | hsa-miR-125b-5p | 0.678 | 0.472 | 0.671 |
| ENSG00000120899 | *PTK2B** | hsa-miR-125b-5p | 0.597 | 0.49 | 0.671 |
| ENSG00000066557 | *LRRC40* | hsa-miR-129-5p | 0.777 | 0.498 | 0.671 |
| ENSG00000108406 | *DHX40* | hsa-miR-501-3p | 1.14 | 0.504 | 0.671 |
| ENSG00000185630 | *PBX1* | hsa-miR-129-5p | 0.928 | 0.512 | 0.671 |

| | | | | | |
|---|---|---|---|---|---|
| ENSG00000163002 | *NUP35* | hsa-miR-129-5p | 1.01 | 0.522 | 0.671 |
| ENSG00000147439 | *BIN3* | hsa-miR-129-5p | 1.01 | 0.533 | 0.671 |
| ENSG00000132600 | *PRMT7* | hsa-miR-138-5p | 0.764 | 0.551 | 0.671 |
| ENSG00000151612 | *ZNF827* | hsa-miR-125b-5p | 0.984 | 0.556 | 0.671 |
| ENSG00000111602 | *TIMELESS* | hsa-miR-195-5p | 1.32 | 0.557 | 0.671 |
| ENSG00000148082 | *SHC3* | hsa-miR-129-5p | 0.579 | 0.57 | 0.671 |
| ENSG00000140848 | *CPNE2* | hsa-miR-501-3p | 1.30 | 0.617 | 0.711 |
| ENSG00000142192 | *APP** | hsa-miR-138-5p, hsa-miR-195-5p | 0.701 | 0.642 | 0.724 |
| ENSG00000163513 | *TGFBR2* | hsa-miR-501-3p | 1.64 | 0.791 | 0.873 |
| ENSG00000128284 | *APOL3* | hsa-miR-125b-5p | 1.54 | 0.82 | 0.887 |
| ENSG00000183117 | *CSMD1* | hsa-miR-195-5p | 0.701 | 0.863 | 0.915 |
| ENSG00000148331 | *ASB6* | hsa-miR-125b-5p | 0.852 | 0.885 | 0.92 |
| ENSG00000090581 | *GNPTG* | hsa-miR-501-3p | 0.913 | 0.974 | 0.991 |
| ENSG00000026950 | *BTN3A1* | hsa-miR-125b-5p | 1.19 | 0.991 | 0.991 |

*Target genes taken from Takousis et al, 2019, all others from TargetScan (see Methods).
‡Only miRNAs analyzed in this study are listed, other miRNAs not studied here may also bind to the same target gene.
FC: fold change between means of gene aggregated normalized TPMs as calculated by sleuth between AD cases and controls.
P-value: derived from gene level likelihood ratio test (LRT) as calculated by sleuth;
Q-value: P-value adjusted using the Benjamini-Hochberg method to control the false discovery rate (FDR) across all 53 genes assessed
Bold font: miRNAs showing nominally significant evidence of differential expression in AD cases vs. controls.
